# Bats use Dip Echolocation to overcome rhythmic noise

**DOI:** 10.64898/2026.08.13.744658

**Authors:** Shivani Hariharan, Susanne S. Babl, Francisco M. López, Nika Jurov, Jochen Triesch, Julio C. Hechavarria

## Abstract

Active sensing systems are known to adapt the structure of sensory signals. Whether they can improve perception by controlling when sensory information is acquired remains unclear. We show that echolocating fruit bats exposed to rhythmic noise preferentially emit calls during recurring low-noise periods, a behaviour we term “dip echolocation”. Dip echolocation occurred in laboratory and wild bats and represents an active-sensing analogue of dip listening in humans. A normative model showed that temporal positioning of calls emerges from a trade-off between sensory information and energetic cost, alongside concurrent adaptations of call structure. Pharmacological inactivation of the frontal auditory field disrupted precise temporal control, implicating a role for frontal cortical circuits in adaptive vocal timing. These findings identify adaptive vocal timing as an active-sensing strategy for overcoming acoustic interference.

## Introduction

Natural acoustic environments are noisy, requiring animals to maintain reliable perception despite external interference (*1–3*). This challenge is especially acute in active sensing systems, where perception depends not only on processing incoming information but also on generating the signals used to probe the environment (*4, 5*). Echolocating bats provide a powerful model for studying this problem because they actively control the acoustic signals that allow perception (5– 10). Studies have shown that bats modify call amplitude, duration, and spectral structure in response to acoustic interference (*2, 6–8*). More recently, work on groups of freely interacting bats has suggested that animals may also adjust the timing of echolocation calls to escape acoustic interference (*9*). Whether this phenomenon reflects a general active-sensing strategy, occurs under natural conditions, confers sensory benefits, and depends on specialized neural mechanisms remains unknown. Such temporal positioning resembles dip listening in humans, where speech perception improves during brief reductions in acoustic masking (*10, 11*). Here, we refer to this alignment of echolocation calls with recurring reductions in interference as dip echolocation (Fig. 1A).

**Fig 1.**
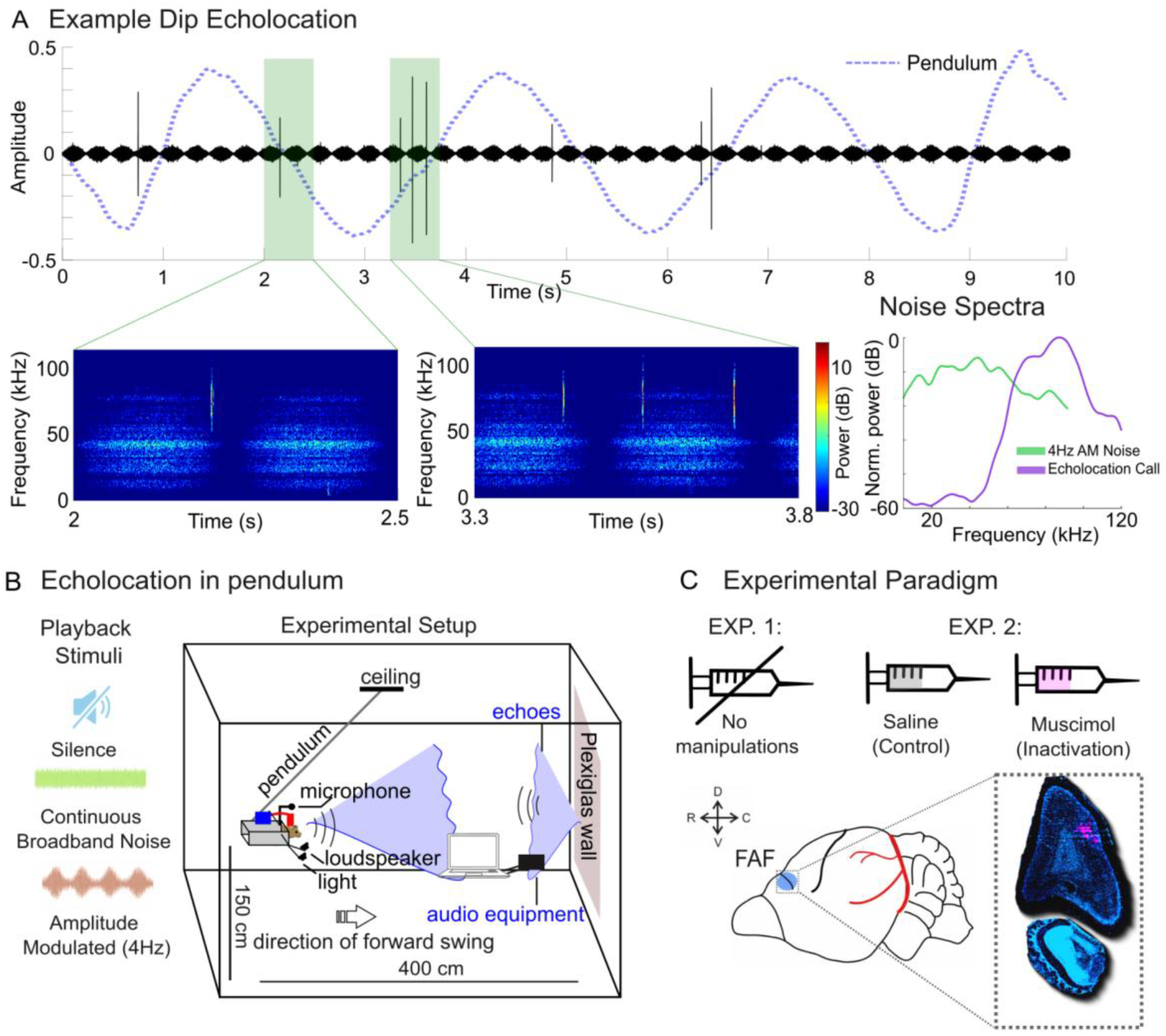
Dip echolocation experimental paradigm. (**A**) Example of dip echolocation. Echolocation calls (gray lines) are aligned to low-amplitude phases of 4 Hz amplitude-modulated (AM) noise (black trace; green shaded regions). The blue dashed trace shows the bat’s position on the pendulum. Corresponding spectrograms of the green shaded areas are shown on the bottom left. Bottom right shows normalized FFT power spectra (dB) of 4 Hz amplitude-modulated (AM) noise and echolocation call. (**B**) Pendulum-based echolocation paradigm. Bats were swung toward a reflective wall while vocalizations were recorded during silence, continuous broadband noise, or 4 Hz amplitude-modulated noise. (**C**) Experimental structure. Experiment 1 measured echolocation behaviour without manipulation. Experiment 2 compared saline controls and muscimol inactivation targeting the frontal auditory field (FAF). Histological verification is shown below.

Understanding dip echolocation requires addressing two fundamental questions. First, what adaptive advantage is gained by controlling when sensory information is acquired? Second, what neural mechanisms allow animals to align sensory actions with favourable moments in the environment? Answering these questions links the functional and mechanistic perspectives that form two of Tinbergen’s classic aims of ethology (*12*). Forebrain circuits contribute to the temporal control of vocal behaviour across a range of species (*13–16*), and the bat frontal auditory field (FAF) has been implicated in vocal-auditory processing and fronto-temporal coordination , making it a candidate substrate for adaptive vocal timing.

To investigate the function and mechanism of dip echolocation in *Carollia perspicillata*, we combined behavioural experiments in the laboratory using a pendulum paradigm with recordings of free-flying bats in the wild. We also performed pharmacological inactivation of the FAF during the pendulum-based echolocation paradigm (Fig. 1B, C). We show that fruit bats preferentially emit echolocation calls during recurring low-noise periods under rhythmic acoustic interference. A normative model indicates that this behaviour emerges from an energy– information trade-off, and FAF inactivation disrupts precise temporal alignment of calls to acoustic dips. Together, these findings suggest that dip echolocation is a behavioural strategy for compensating for acoustic interference.

## Results

### Bats adapt call structure and timing under acoustic interference

To determine how bats adapt echolocation behaviour under acoustic interference, we exposed bats to periods of silence, continuous broadband noise (CN), and 4 Hz amplitude-modulated (AM) noise, while the animals were swung on a pendulum simulating a controlled flight path (Fig. 1B). We then measured call amplitude, duration, spectral structure, and timing of the emitted echolocation calls (Fig. 2A–I). While previous studies have largely examined these vocal adaptations individually, we quantified multiple parameters within the same trials to investigate not only how bats compensate for acoustic interference, but also whether predictable dips in noise enable them to strategically time sensory information acquisition.

**Fig 2.**
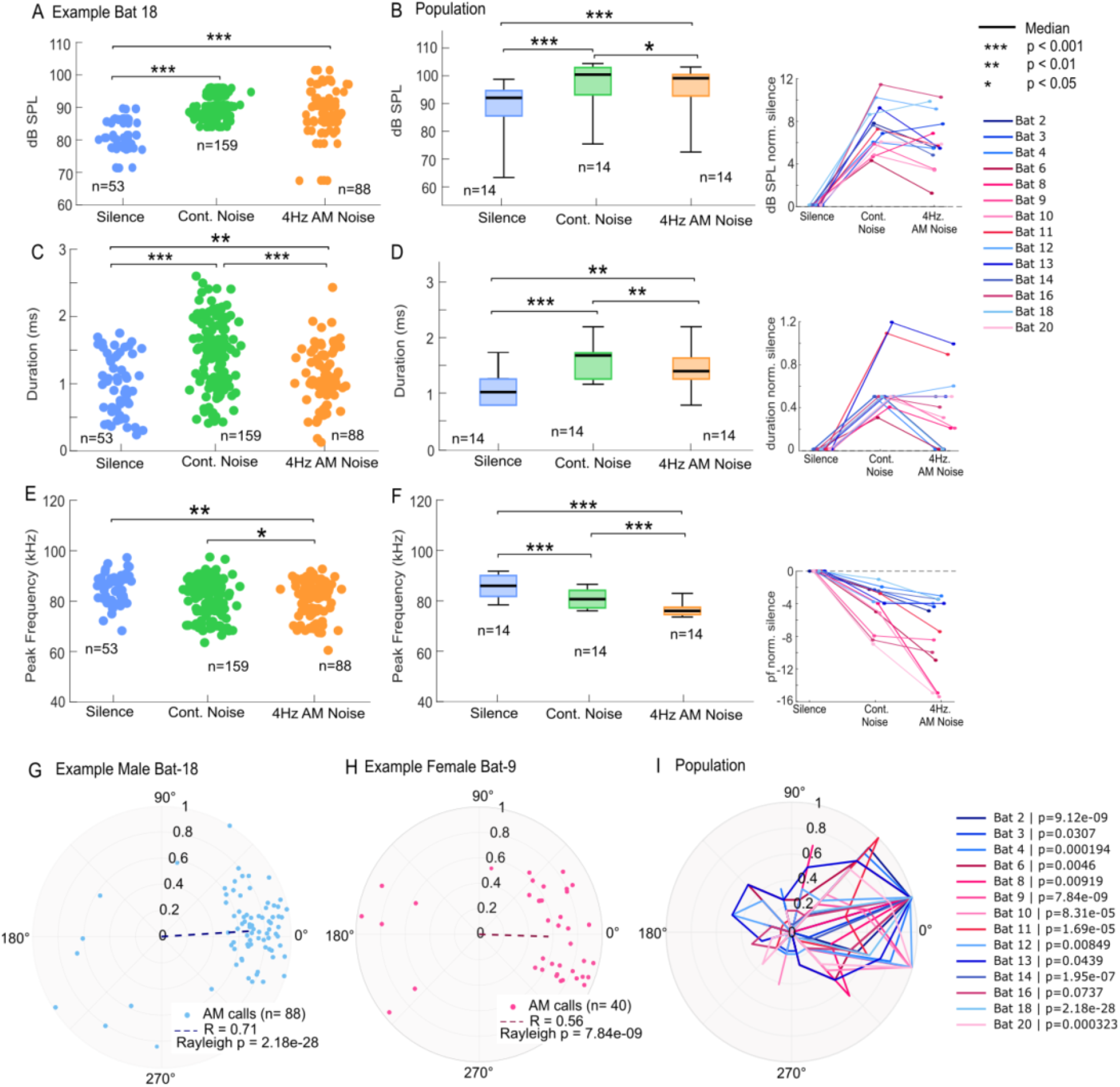
Echolocation responses to acoustic interference. (**A–F**) Call sound pressure level (SPL), duration, and peak frequency in silence, continuous broadband noise, and amplitude-modulated (AM) noise. (**A, C, E**) Individual calls from an example bat (Wilcoxon rank-sum tests). (**B, D, F**) Population distributions (n = 14 bats; black lines denote medians, Wilcoxon signed-rank tests). Median call parameters of individual bats are shown on the right. Acoustic interference increased call SPL, increased call duration, and lowered peak frequency across conditions (*P* < 0.05 to *P* < 0.001, as indicated). Exact statistical results, including p-values and Cliff’s delta effect sizes, for Example bat 18 and population analyses are provided in Supplementary Tables S2a and S2b, respectively. (**G–I**) In AM noise, calls were phase locked to the modulation cycle. (**G, H**) Example bats showing significant non-uniform phase distributions. (**I**) Population phase distributions demonstrating significant phase locking in 13 of 14 bats (Rayleigh test, *P* ≤ 0.0439).

Bats responded to acoustic interference through concurrent changes in multiple vocal parameters. Relative to silence, both continuous and AM noise elicited increases in call amplitude of ∼5–10 dB (Fig. 2A,B; Wilcoxon rank-sum tests (individual), Wilcoxon signed-rank tests (population): all *P* < 0.001, Cliff’s delta (d) = 0.54–0.86), consistent with a Lombard-like response (*2, 3, 19*). Call duration likewise increased under both noise conditions (Fig. 2C, D; all P ≤ 0.006, Cliff’s delta (d) = 0.27–0.76). Peak frequency generally decreased under acoustic interference (Fig. 2E, F). At the population level, all comparisons were significant (all *P* ≤ 0.00024, Cliff’s delta (d) = −0.92 to −0.62). These effects were evident at both the individual and population levels (n = 14 bats; 7 males, 7 females) and occurred under both continuous and rhythmic noise, indicating that bats employ a coordinated response to acoustic interference rather than relying on compensatory adjustment of a single parameter.

Among all vocal adaptations, the strongest effect emerged in the temporal domain. Under 4-Hz AM noise, bats phase locked call timing to the modulation cycle and preferentially emitted calls during low-noise phases (Fig. 2G–I). Phase locking was significant in 13 of 14 bats (Rayleigh tests, *P* ≤ 0.0439) and occurred in both sexes. Bats therefore responded to rhythmic interference not only by modifying call structure but also by strategically timing when they gathered information from the environment to periods of reduced masking.

### Wild Bats exhibit dip echolocation

A key question was whether bats use dip echolocation as part of their natural behaviour, rather than as an adaptation specific to laboratory conditions. To address this issue, we recorded vocalizations of wild *Carollia* bat populations in the neotropical rainforest in Panama, while playing AM noise through a speaker in three behavioural contexts: perching, free flight within a flight cage, and approaches to an outside feeding station (Fig. 3A).

**Fig 3.**
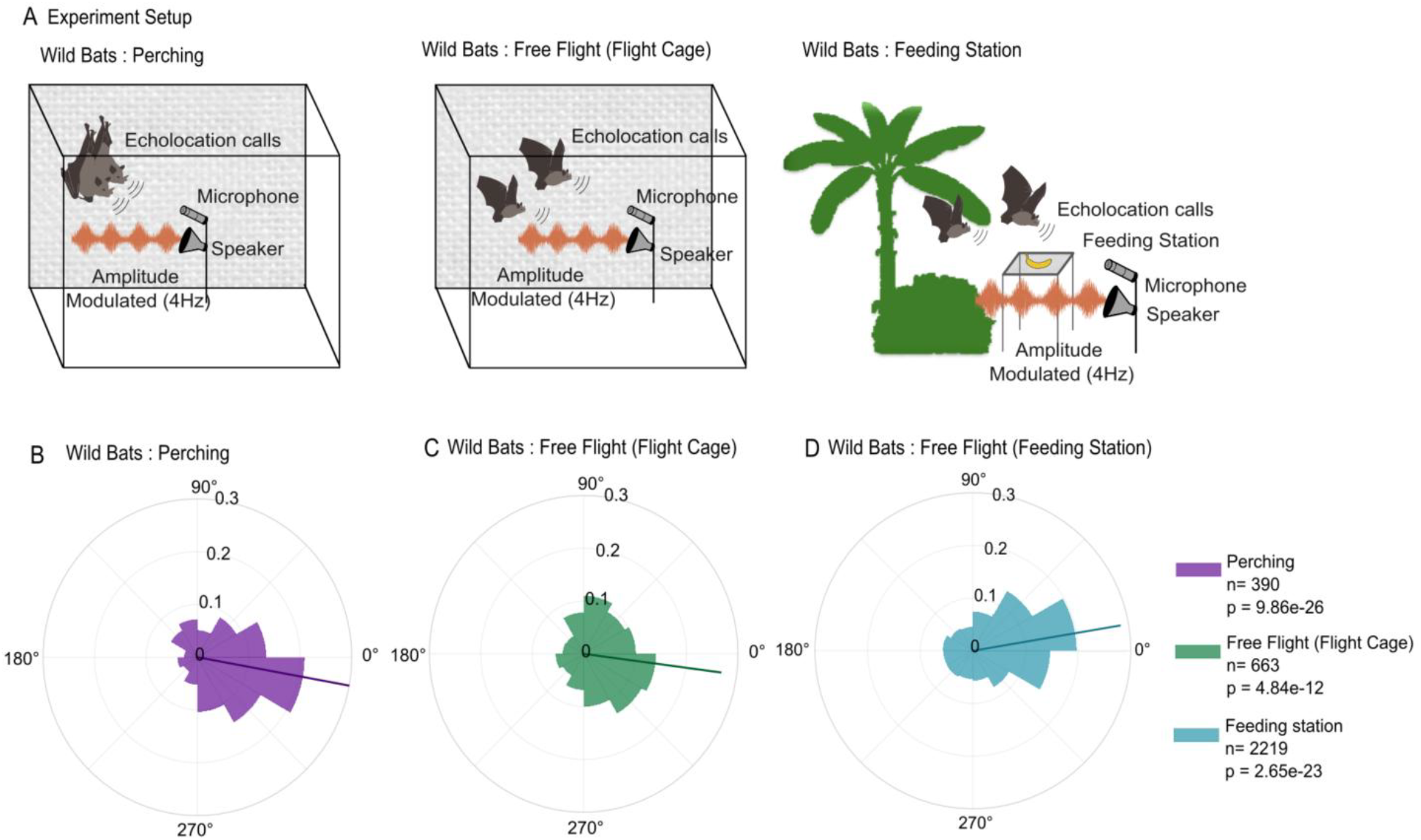
Dip echolocation across behavioural contexts in the wild. (**A**) Experimental setup with acoustic playback. **(B–D**) Wild bats timed calls non-uniformly relative to the AM cycle during perching, free flight in a flight cage, and approaches to a feeding station (Rayleigh tests, *P* < 0.001).

Wild bats exhibited adaptive vocal timing in all contexts examined. During perching, echolocation calls were non-uniformly distributed relative to the modulation cycle and preferentially occurred during low-noise phases (Rayleigh test: *P* = 9.86 × 10^−26^; Fig. 3B). The same temporal strategy was evident during free flight within a flight cage (Rayleigh test: *P* = 4.84 × 10^−12^; Fig. 3C) and during approaches to a feeding station (Rayleigh test: *P* = 2.65× 10^−23^; Fig. 3D). Bats consistently positioned echolocation calls within low noise phases under rhythmic acoustic interference across behavioural states ranging from stationary perching to active foraging. The similarity between laboratory and field observations indicates that dip echolocation is a naturally expressed active-sensing strategy and suggests a robust solution to rhythmic acoustic interference.

### Normative modelling predicts adaptive call timing

Having established dip echolocation as a naturally expressed active-sensing strategy, we next asked why dip echolocation should emerge under acoustic interference. We developed a normative model in which echolocation behaviour maximizes utility defined by a trade-off between sensory information gain and energetic cost (Fig. 4A) (*20*). In the model, call amplitude and duration increase echo reliability but require greater energetic investment (*9, 21, 22*), whereas call frequency and timing influence the degree of overlap between echolocation calls and masking noise (see Materials and Methods and Supplementary Table S1). We evaluated the model under the same acoustic conditions used experimentally: silence, continuous broadband noise, and 4-Hz amplitude-modulated (AM) noise.

**Fig 4.**
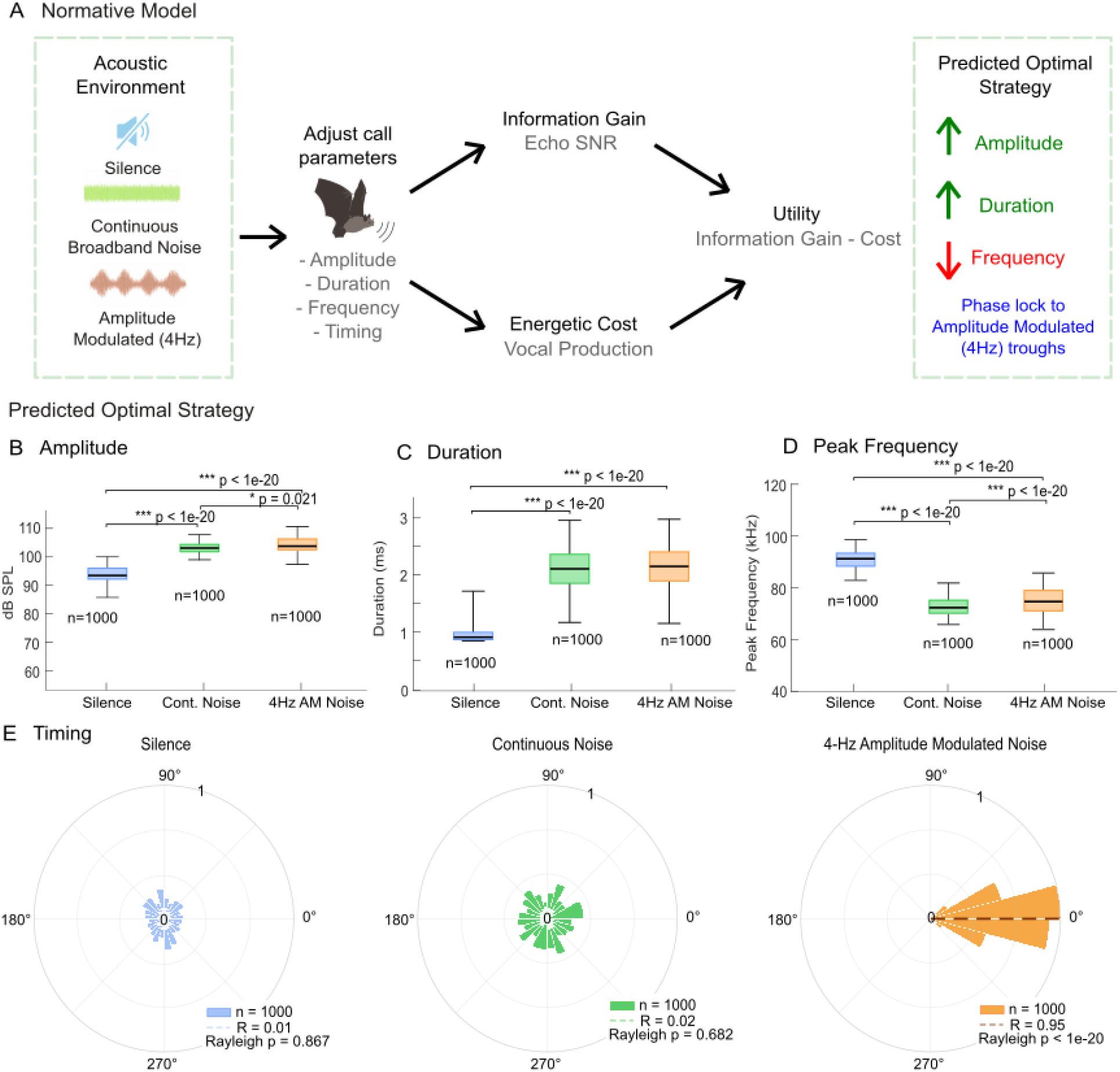
Normative model predictions for call adaptation under acoustic interference. (**A**) Schematic of the normative model. Call parameters (amplitude, duration, frequency, and timing) are adjusted to maximize utility, defined as the trade-off between information gain (improved echo signal-to-noise ratio) and energetic cost of vocal production. The model predicts that animals should increase call amplitude and duration in noisy environments while decreasing call frequency. In temporally structured maskers, call timing is predicted to align with low-noise periods to maximize utility. (**B–D**) Predicted call amplitude (SPL), duration, and peak frequency in silence, continuous broadband noise, and AM noise (n = 1000 simulated calls per condition). The model predicts increased amplitude and duration, together with decreased peak frequency, under acoustic interference. (**E**) Predicted call timing relative to the modulation cycle. Calls are uniformly distributed in silence and continuous noise but become strongly phase locked in 4-Hz AM noise, with calls concentrated near modulation troughs where masking is minimized (Rayleigh test, *P* ≤ 10−20, R = 0.95).

The model predicted that acoustic interference should favour increases in call amplitude and duration and, when noise overlapped with the bat’s call frequency, a reduction in peak frequency (Fig. 4B–D). Under AM noise, the model further predicted that bats should preferentially emit calls during low-noise phases of the modulation cycle, where echo reliability is maximized (Fig. 4E). This indicates that dip echolocation emerges naturally from the energy–information trade-off and reproduces the major behavioural adaptations observed experimentally. These findings suggest that temporal positioning of acoustic signals can improve sensory performance without additional energetic expenditure, making vocal timing an efficient strategy for active sensing under interference.

### Frontal cortex inactivation disrupts dip echolocation

This raised the question of whether dip echolocation depends on higher-order cortical processing. Because the behaviour requires aligning vocal output to the temporal structure of ongoing acoustic interference, we hypothesized that frontal circuits contribute to its implementation, similar to their roles in timing-dependent vocal behaviours in birds, singing mice, and marmosets (*10, 11, 13–16, 23, 24*). To test this, we pharmacologically inactivated the frontal auditory field (FAF) using muscimol and compared echolocation behaviour to a saline condition within the same animals (Fig. 5A–L). Histological analyses confirmed that muscimol diffusion remained confined to the targeted region (layer V) (Fig. 1C).

**Fig 5.**
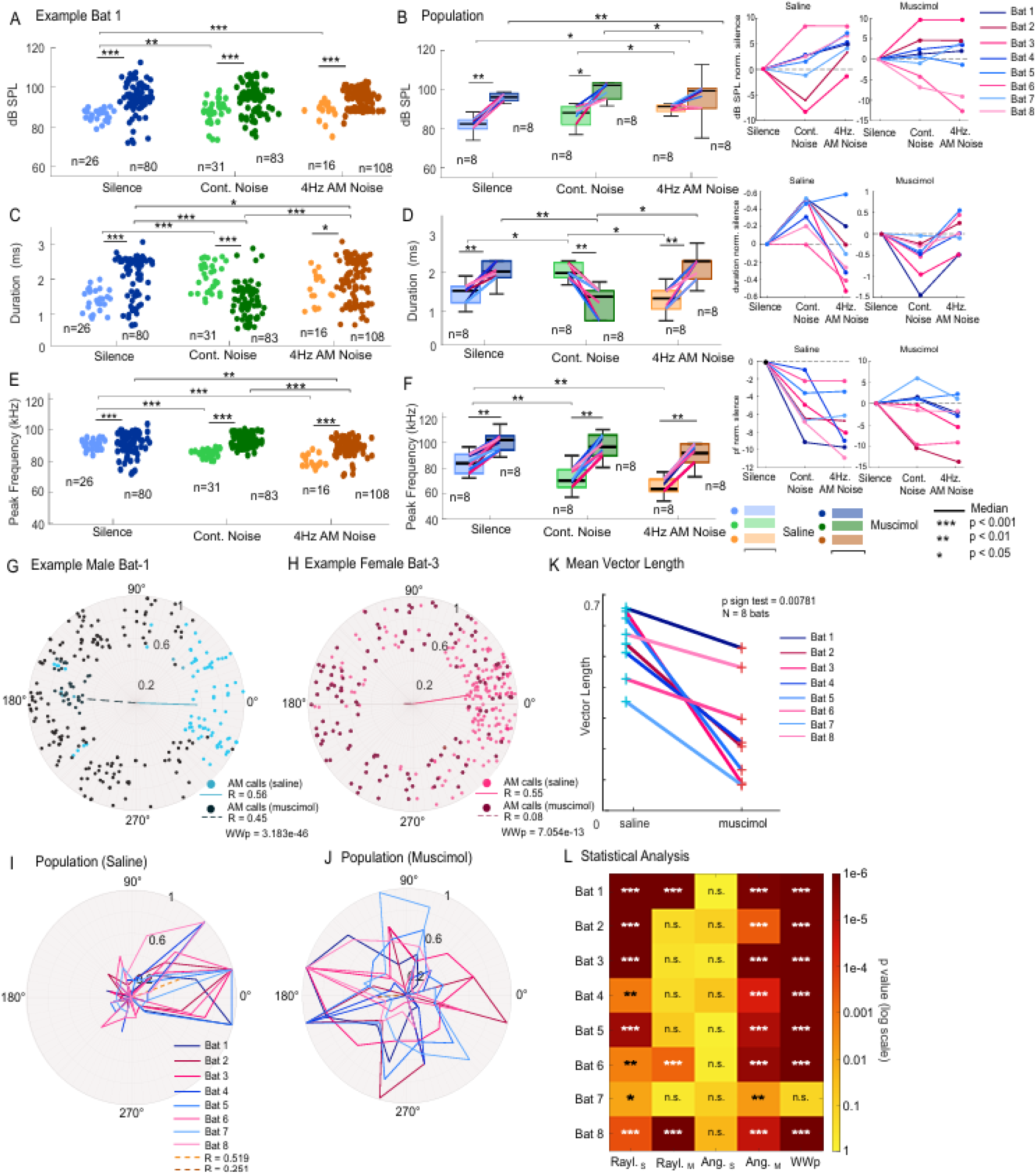
Inactivating FAF alters call structure and disrupts temporal coordination. (A–F) Call SPL, duration, and peak frequency in silence, continuous broadband noise, and AM noise under saline and muscimol conditions. (A, C, E) Individual calls from an Example bat. (B, D, F) Population distributions (n = 8 bats; black lines denote medians). Muscimol altered call level, duration, and peak frequency across acoustic conditions (*exact statistical results for Example bat 1 and population analyses are provided in Supplementary Tables S3a and S3b, respectively*). (G–I) In AM noise, muscimol reduced phase locking to the modulation cycle. (G, H) Example bats timing calls under saline and muscimol. (I, J) Population phase distributions under saline and muscimol. Phase locking was stronger under saline than after muscimol. (K) Mean vector length decreased in all eight bats after muscimol (sign test, P = 0.00781). (L) Statistical summary across bats; stars indicate Rayleigh and Watson–Williams test significance (*\* P < 0*.*05, ** P < 0*.*01, *** P < 0*.*001*). Exact p-values are reported in Supplementary Table S4.

Bats continued to produce echolocation calls following FAF inactivation, indicating that cortical input is not required for call generation itself. Call amplitude increased under muscimol across conditions by 6.5 ± 5.5 dB (mean ± SD, n =8) (Fig. 5A, B, *see Supplementary Tables S3a and S3b for statistical values*), although bats retained the ability to increase call level under acoustic interference. Call duration and peak frequency also changed following muscimol application relative to saline controls (Fig. 5C–F).

The strongest effect of FAF inactivation emerged in the temporal domain. Under saline conditions, all bats exhibited robust dip echolocation, i.e., they preferentially emitted echolocation calls during low-noise phases of the modulation cycle (Fig. 5G–J). Following muscimol application, phase locking weakened, calls were distributed more broadly across noise phases, and preferred call timing frequently shifted away from acoustic dips. Mean resultant vector length decreased in every animal (sign test: *P* = 0.00781, n = 8 bats; Fig. 5K), despite a reduction in call rate under muscimol (Supplementary Fig. S1). Whereas all bats showed significant phase clustering under saline conditions (Rayleigh tests: all *P* ≤ 0.010), only 3 of 8 bats remained significantly phase locked after FAF inactivation (Fig. 5L). Notably, the three bats that remained significantly phase locked showed substantial shifts in preferred phase under muscimol. FAF inactivation impaired the temporal coordination of echolocation behaviour, consistent with a role for frontal cortical circuits in adaptive vocal timing (*17, 18, 25, 26*).

In summary, these results identify the FAF as a cortical contributor to adaptive vocal timing under acoustic interference. Although FAF inactivation altered multiple call parameters, its most consistent effect was a disruption of dip echolocation, indicating that frontal cortical circuits contribute to aligning calls with favourable moments in the acoustic environment. These findings suggest that, in bats, temporal optimization of active sensing depends, at least in part, on higher-order cortical processing.

## Discussion

Although previous work has shown that bats can adjust vocal timing to avoid acoustic interference during social interactions and that this ability disappears when noise modulation rates exceed ∼20Hz (*9*), it remained unclear whether such temporal control reflects a general principle of active sensing or a context-dependent response to vocal interactions. By combining laboratory experiments, field observations, normative modelling, and cortical manipulation, we establish dip echolocation as a naturally employed strategy for overcoming rhythmic acoustic interference and identifying its adaptive value.

Exploiting windows of reduced acoustic interference in masking noise appears to be a widespread solution to sensory interference. Similar benefits have been described in humans listening to speech in fluctuating maskers, where brief reductions in masker level improve speech perception through “listening in the dips” (*10, 11*). These parallels suggest that bats and humans may converge on related strategies for improving perception under acoustic interference despite relying on very different sensory systems.

The normative model provides a potential explanation for why precise vocal timing emerges under rhythmic acoustic interference. Timing occurred as a particularly efficient control variable because sensory information could be increased by exploiting transient windows in interference without requiring additional energetic investment. Dip echolocation therefore arose naturally from an energy–information trade-off (*20, 21*), suggesting that temporal positioning of sensory signals may represent a general strategy for improving active sensing under fluctuating environmental conditions.

Frontal inactivation disrupted the temporal alignment of echolocation calls to acoustic dips while preserving vocal production, suggesting that fronto-auditory circuits contribute to adaptive vocal timing in mammals. A related phenomenon, termed dip listening, has been described in tree frogs, where receivers improve signal detection by listening during brief dips in background acoustic interference (*27*). Comparable behavioural solutions that have evolved in animals lacking a neocortex suggest that exploiting predictable acoustic dips represents a general solution to sensory interference, implemented by different neural circuits across vertebrates. Our findings implicate the frontal cortex as a critical region of the neural network that supports bats in actively exploiting predictable acoustic dips during echolocation. Together, the behavioural, modelling, and inactivation results link the adaptive value and mechanistic basis of dip echolocation, addressing two central aims of ethology: understanding both the function of behaviour and the mechanisms that generate it. (*12*)

Our findings suggest that controlling when sensory information is acquired may be as important as controlling the properties of the signals used to acquire it, thus establishing the principle of dip sensing as a central strategy of adaptive sensing across species and sensory modalities.

## Acknowledgments

We thank Gisa Prange and the veterinary team at the Institute for Cell Biology and Neuroscience at Goethe University Frankfurt for their support. We thank the Smithsonian Tropical Research Institute, especially Rachel Page and Gregg Cohen, for providing access to facilities and logistical support. This research was supported by the German Research Foundation (Deutsche Forschungsgemeinschaft, DFG) (SPP 2411 - “Sensing LOOPS”)

## Author contributions

J.C.H. and J.T. conceived the study and supervised the project. S.H. collected and analysed the experimental data from the laboratory studies under the supervision of J.C.H. S.S.B. conducted the field studies in Panama, S.H. analysed the field data. F.L. developed the normative model and, together with N.J. and J.T., performed the computational modelling and analyses. S.H. and J.C.H. wrote the first draft of the manuscript. All authors contributed to interpretation of the results, revised the manuscript, and approved the final version.

## References

1. H. Brumm, S. A. Zollinger, The evolution of the Lombard effect: 100 years of psychoacoustic research. Behaviour 148, 1173–1198 (2011).

2. J. Luo, S. R. Hage, C. F. Moss, The Lombard Effect: From Acoustics to Neural Mechanisms. Trends in Neurosciences 41, 938–949 (2018).

3. S. A. Zollinger, H. Brumm, The Lombard effect. Curr Biol 21, R614–615 (2011).

4. M. E. Nelson, M. A. MacIver, Sensory acquisition in active sensing systems. J Comp Physiol A Neuroethol Sens Neural Behav Physiol 192, 573–586 (2006).

5. C. F. Moss, A. Surlykke, Probing the Natural Scene by Echolocation in Bats. Front. Behav. Neurosci. 4 (2010).

6. S. R. Hage, W. Metzner, Potential effects of anthropogenic noise on echolocation behaviour in horseshoe bats. Communicative & Integrative Biology 6, e24753 (2013).

7. E. Amichai, G. Blumrosen, Y. Yovel, Calling louder and longer: how bats use biosonar under severe acoustic interference from other bats. Proc Biol Sci 282, 20152064 (2015).

8. E. H. Gillam, N. Ulanovsky, G. F. McCracken, Rapid jamming avoidance in biosonar. Proc Biol Sci 274, 651–660 (2007).

9. A. Kiai, J. Clemens, M. Kössl, D. Poeppel, J. Hechavarría, Flexible control of vocal timing in Carollia perspicillata bats enables escape from acoustic interference. Commun Biol 6, 1153 (2023).

10. J. M. Festen, Contributions of comodulation masking release and temporal resolution to the speech-reception threshold masked by an interfering voice. J Acoust Soc Am 94, 1295–1300 (1993).

11. K. K. Jensen, J. G. W. Bernstein, The fluctuating masker benefit for normal-hearing and hearing-impaired listeners with equal audibility at a fixed signal-to-noise ratio. J Acoust Soc Am 145, 2113–2125 (2019).

12. N. Tinbergen, On aims and methods of Ethology. Zeitschrift für Tierpsychologie 20, 410–433 (1963).

13. J. I. Benichov, S. E. Benezra, D. Vallentin, E. Globerson, M. A. Long, O. Tchernichovski, The Forebrain Song System Mediates Predictive Call Timing in Female and Male Zebra Finches. Curr Biol 26, 309–318 (2016).

14. J. I. Benichov, D. Vallentin, Inhibition within a premotor circuit controls the timing of vocal turn-taking in zebra finches. Nat Commun 11, 221 (2020).

15. D. E. Okobi, A. Banerjee, A. M. M. Matheson, S. M. Phelps, M. A. Long, Motor cortical control of vocal interaction in neotropical singing mice. Science 363, 983–988 (2019).

16. D. Dohmen, S. R. Hage, Limited capabilities for condition-dependent modulation of vocal turn-taking behaviour in marmoset monkeys. Behavioural Neuroscience 133, 320–328 (2019).

17. F. García-Rosales, L. López-Jury, E. González-Palomares, J. Wetekam, Y. Cabral-Calderín, A. Kiai, M. Kössl, J. C. Hechavarría, Echolocation-related reversal of information flow in a cortical vocalization network. Nat Commun 13, 3642 (2022).

18. F. García-Rosales, L. López-Jury, E. González-Palomares, Y. Cabral-Calderín, J. C. Hechavarría, Fronto-Temporal Coupling Dynamics During Spontaneous Activity and Auditory Processing in the Bat Carollia perspicillata. Front Syst Neurosci 14, 14 (2020).

19. J. Luo, H. R. Goerlitz, H. Brumm, L. Wiegrebe, Linking the sender to the receiver: vocal adjustments by bats to maintain signal detection in noise. Sci Rep 5, 18556 (2015).

20. P. Sommer, F. Zeldenrust, P. Jedlicka, A. D. Bird, J. Triesch, Neuronal degeneracy reflects context-specific trade-offs between energy and information. bioRxiv [Preprint] (2026). 10.64898/2026.03.02.708507.

21. F. M. López, B. E. Shi, J. Triesch, “Chapter Five - Efficient coding in active perception: A developmental perspective on autonomous control” in Computational and Modeling Approaches to Development, A. S. Warlaumont, J. J. Lockman, Eds. vol. 70 of Advances in Child Development and Behaviour, pp. 117–156.

22. C. Chiu, W. Xian, C. F. Moss, Adaptive echolocation behaviour in bats for the analysis of auditory scenes. J Exp Biol 212, 1392–1404 (2009).

23. A. Banerjee, D. Vallentin, Convergent behavioural strategies and neural computations during vocal turn-taking across diverse species. Current Opinion in Neurobiology 73, 102529 (2022).

24. L. M. Martin, F. García-Rosales, M. J. Beetz, J. C. Hechavarría, Processing of temporally patterned sounds in the auditory cortex of Seba’s short-tailed bat,Carollia perspicillata. European Journal of Neuroscience 46, 2365–2379 (2017).

25. A. Eiermann, K. H. Esser, Auditory responses from the frontal cortex in the short-tailed fruit bat Carollia perspicillata. Neuroreport 11, 421–425 (2000).

26. J. S. Kanwal, M. Gordon, J. P. Peng, K. Heinz-Esser, Auditory responses from the frontal cortex in the mustached bat, Pteronotus parnellii. Neuroreport 11, 367–372 (2000).

27. A. Vélez, M. A. Bee, Dip listening and the cocktail party problem in grey treefrogs: signal recognition in temporally fluctuating noise. Animal Behaviour 82, 1319–1327 (2011).

