## Supplementary Files for "Bats use Dip Echolocation to overcome rhythmic noise"

**The PDF file includes:**

Materials and Methods
Fig. S1
Tables S1-S4
References

### Materials and Methods

Animals

For laboratory experiments, we used 22 adult bats (*Carollia perspicillata)*. Fourteen animals (7 males, 7 females) participated in Experiment 1 and did not undergo any experimental manipulation. An additional eight bats (4 males, 4 females) participated in Experiment 2, which involved pharmacological inactivation of the frontal auditory field (FAF).

These animals were obtained from the breeding colony at the Institute for Cell Biology and Neuroscience, Goethe University Frankfurt, Germany. We housed bats in a temperature-controlled room (28°C, ~60% humidity). All procedures complied with German regulations governing animal experimentation and were approved by the *Regierungspräsidium Darmstadt (permits no. F104/57, FU-1126 /FR2007).*

Experiment 1

We positioned bats on a platform attached to the mass of a pendulum apparatus. A foam layer between the acrylic plate and the platform ensured stability and minimized movement. We gently positioned each bat between foam layers and secured an acrylic plate above the animal to maintain a stable posture.

We manually released the pendulum, allowing the bat to swing toward a large acrylic wall (50 × 150 cm). We recorded echolocation calls continuously for 10 s during pendulum motion, including both forward and backward swings. During this period, bats emitted echolocation pulses and received returning echoes from the wall.

We recorded ultrasonic vocalizations using a condenser microphone (CM16/CMPA, Avisoft Bioacoustics, Germany) mounted on the pendulum mass above the bat's head. We positioned the microphone membrane approximately 4 cm from the bat's ears. After recording, we manually stopped the pendulum.

Experiment 2

We used the same behavioural procedure as in Experiment 1 but added pharmacological inactivation of the FAF. We anesthetized animals with ketamine (*10 mg kg*$^{-1}$*, Ketavet,* Pfizer) and xylazine (*38 mg kg*$^{-1}$*, Rompun, Bayer*). We applied a local anesthetic (*ropivacaine hydrochloride, 2 mg ml*$^{-1}$*; Fresenius Kabi, Germany*) subcutaneously around the scalp before incision. We then made a midline incision to expose the skull and performed small bilateral craniotomies above the FAF. After surgery, we allowed animals to recover for at least 2 days before behavioural testing.

On the day of the experiment, we applied either sterile saline (0.9%; control) or muscimol (8 mM in 0.9% saline) topically to the cortical surface of the FAF. We applied drug solutions in 1 µl aliquots and began behavioural testing 30 min after application to allow for cortical inactivation. Experimental sessions did not exceed 4 h per day. We provided water every 1–1.5 h and terminated experiments if animals showed signs of distress.

Acoustic Stimulation

We generated acoustic stimuli in *MATLAB (R2024b, MathWorks)* and presented them using Avisoft-RECORDER USGH (version 4.3.00). We used three stimulus conditions in both experiments: silence, continuous broadband noise (5–90 kHz), and amplitude-modulated (AM) broadband noise (5–90 kHz).

We modulated the AM stimulus at 4 Hz, based on previous work characterizing the temporal structure of spontaneous *Carollia* vocalizations (*1*, *2*). We presented stimuli for 10s during pendulum motion and repeated each condition five times per session in randomized order.

We delivered acoustic stimuli through a directional speaker positioned approximately 10 cm from the bat. Audio signals were routed through a Fireface 400 sound card (RME) and amplifier. Stimulus intensity was controlled using 30 dB attenuation. Based on the speaker calibration, playback stimuli were presented at approximately 50–60 dB SPL RMS at the bat's position.

The spectral range of the noise stimuli partially overlapped with the frequency range of *C. perspicillata* echolocation calls. Bats produced echolocation pulses naturally during pendulum motion, and we recorded these calls together with their returning echoes.

Audio and video recordings

We recorded ultrasonic vocalizations using an Avisoft CM16 condenser microphone (16-bit resolution, 250-kHz sampling rate) connected to an UltraSoundGate 116Hm interface (Avisoft Bioacoustics).

We recorded videos using a Basler AG camera, positioned above the speaker and outside the direct acoustic path. Infrared illumination below the bat enabled recording in darkness.

We synchronized audio and video using two trigger channels. One TTL pulse triggered acoustic recording via the UltraSoundGate interface. A second trigger (RP2 TDT system) activated an LED visible to the camera, enabling precise offline alignment of audio and video data.

Field experiments

Playback experiments in wild *Carollia* were performed at the Smithsonian Tropical Research Institute in Gamboa, Panama. All field work was approved by Panamanian authorities (*Ministerio de Ambiente de Panama,* permit no. ARG-0094-2025*)* and conducted according to permits of the Smithsonian Tropical Research Institute Animal Care and Use Committee issued to Rachel Page (permit no. SI-23001-29).

For experiments in the flight cage, two adult *Carollia perspicillata* (one male, one nonreproductive female) were captured outside their roost using mist nets in November 2025. The bats were housed together in a mesh-lined cage (30 x 50 x 50 cm), with food (banana) and water provided ad libitum, and were released at the site of capture after four days.

For vocal recordings during 4 Hz AM noise, both bats were released together into a flight cage (5 x 5 m) at dusk and allowed to adapt to the cage for >1 h. Once bats were perched on a wall of the cage, a speaker and microphone were positioned approximately 1 m below them. AM noise was played in 30 second bouts interleaved with two-minute periods of silence for a total duration of 15 minutes, while vocalizations were recorded using an Avisoft CM16 condenser microphone connected to an UltraSoundGate 116Hm interface and an Avisoft ultrasonic Speaker Vifa connected to an UltraSoundGate 116H player. To obtain recordings during flight, the same playback protocol was used, but perched bats were occasionally approached by the experimenter to induce short flights within the cage.

To assess whether bats used dip echolocation during natural foraging behavior, a feeding station baited with banana was established in an open area at the edge of the rainforest. Use of the station by *Carollia spp.* was confirmed for several days prior to the experiment using camera recordings and acoustic monitoring. For playback experiments, a speaker and microphone were positioned at 0.5 m distance facing the feeding station. During an active bat visitation period, AM noise was presented in 30 second bouts interleaved with one-minute periods of silence for approximately 20 minutes in total.

Acoustic analysis

We processed acoustic recordings in Avisoft SASLab Pro (Avisoft Bioacoustics). We reduced background noise using the built-in noise reduction function (FFT length: 256; precision: 16), which attenuates low-amplitude spectral components while preserving higher-amplitude signals. Echolocation calls and echoes exceeded this threshold and were unaffected.

We manually labeled echolocation calls based on characteristic spectral features of *C. perspicillata* (*3*, *4*). We then extracted acoustic parameters using the Automatic Parameter Measurement function in Avisoft SASLab Pro, including call duration, peak frequency, and sound pressure level (dB SPL).

Phase-locking analysis

For temporal analyses, we filtered recordings using a finite impulse response (FIR) band-pass filter (40–60 kHz) implemented in Avisoft SASLab Pro. We applied the filter using a Hamming window.

We analyzed filtered audio files and call onset timestamps using custom MATLAB scripts. We computed the amplitude envelope of the stimulus and extracted its instantaneous phase using the Hilbert transform. We then sampled phase values at call onset times. We pooled phase values across calls and animals to compute circular statistics, including mean phase direction and resultant vector length. We assessed non-uniformity using the Rayleigh test.

Statistical analysis

We performed all statistical analyses in *MATLAB (R2024b)* using custom scripts. For each bat and condition, we extracted call intensity (dB SPL), duration (ms), and peak frequency (kHz). For representative animals, we compared call-level distributions using two-sided Wilcoxon rank-sum tests. For population-level analyses, we calculated a single median value per bat and condition and performed paired comparisons using two-sided Wilcoxon signed-rank tests. Effect sizes were quantified using Cliff's delta (d), with values of ~0.2, ~0.5, and ≥0.8 interpreted as small, medium, and large effects, respectively. Positive effect sizes indicate larger values in the second-listed condition relative to the first-listed condition. In Experiment 1, we compared silence, broadband and amplitude-modulated noise conditions. In Experiment 2, we extended this design by additionally comparing saline and muscimol conditions within each stimulus context.

We visualized population-level distributions using kernel density–based violin plots with paired observations connected.

For circular analyses, we used the *CircStat* toolbox in MATLAB. We computed the instantaneous phase of the amplitude-modulated stimulus using the Hilbert transform and extracted phase values at call onset.

For each bat and condition, we assessed deviations from uniformity using the Rayleigh test. We report the mean phase angle for each condition in degrees, with clockwise and counterclockwise directions indicated and compared mean phase angles between saline and muscimol conditions using the Watson–Williams test.

Statistical significance was defined as $P < 0.05$ (*), $P < 0.01$ (**), and $P < 0.001$ (***).

Normative Model

We developed a minimal normative model to study whether the observed bat behaviors were consistent with an optimal adaptation strategy under acoustic interference (*1, 5–7*). The model assumes that each echolocation pulse is defined by four parameters controlled by the bat: amplitude A, duration d, peak frequency f, and phase $\phi$. For each condition (silence, constant broadband noise, and 4 Hz AM noise), the optimal adaptation was defined as the set of parameters that maximize a utility function balancing sensory information gain against energetic cost (*8, 9*).

*Information gain*

We modeled the information gain of an echolocation call as a function of the signal-to-noise ratio (SNR) of the returning echo. The signal strength was assumed to increase exclusively with acoustic power and call duration (*8*)

$$S(A,d) = P(A)\,.d$$

where P(A) denotes relative acoustic power. To express amplitude in dB SPL, we converted it to acoustic power relative to a reference amplitude $A$ref as

$$P(A) = 10^{\frac{(A - Aref)}{10}}$$

To model the level of noise relative to the signal, we assume that bats can filter the incoming auditory inputs with respect to the properties of their calls. That is, rather than processing to the entire input, bats can selectively filter or attend to the relevant frequency and temporal components (9). The effective noise level was modeled as

$$N(f, \phi) = \sigma^2 + N_f(f)\, N_\phi(\phi)$$

where $\sigma 2$ is a baseline noise level in the environment, $N_f(f)$ captures frequency-dependent noise, and $N_\phi(\phi)$ captures phase-dependent noise in the AM condition

The resulting SNR was

$$SNR(A, f, d, \phi) = \frac{S(A, d)}{N(f, \phi)}$$

Information gain was then defined as a sigmoid function of SNR:

$$I(A, f, d, \phi) = \frac{1}{1 + exp\left[-\beta(SNR(A, f, d, \phi) - \theta)\right]}$$

where $\theta$ determines the inflection point at which information gain transitions from low to high and $\beta$ controls the slope of this transition.

*Noise conditions*

In the silence condition, the frequency-dependent noise term was set to zero

$$N_f(f) = 0$$

Under acoustic interference, we modeled spectral noise as a Gaussian function:

$$N_f(f) = N_B exp\left[-\frac{\left(f - f_n\right)^2}{2\sigma_f^{\,2}}\right]$$

where $N_B$ controls the magnitude of noise, $f_n$ the center of the Gaussian function and $\sigma f$ its bandwidth. For silence and continuous broadband noise, the temporal noise component was constant:

$$N_{\phi}(\phi) = 1$$

For AM noise, temporal masking varies sinusoidally with phase:

$$N_{\phi}(\phi) = N_{min} + \frac{1}{2}\left(N_{max} - N_{min}\right)\left[1 - \cos\left(\phi - \phi_{m}\right)\right]$$

where $N_{\min}$ and $N_{\max}$ define the trough and peak noise levels, and $\phi m$ defines the phase of minimum noise. Thus, the effective noise was lowest at the dip of the AM cycle.

*Energetic cost*

The energetic cost of producing a call was modeled as the sum of independent contributions of each call parameter:

$$C(A, f, d) = \lambda_{c} + \lambda_{A} C_{A}(A) + \lambda_{d} C_{d}(d) + \lambda_{f} C_{f}(f)$$

Where $\lambda c$ is a baseline cost, $\lambda_A$, $\lambda_d$, and $\lambda_f$ control the cost of call amplitude, duration, and frequency, respectively. We assume that the phase does not change the energetic cost of a call. For amplitude and duration, the cost functions were directly set to the signal strength functions:

$$C_{A}(A) = P(A), \;\; C_{d}(d) = d$$

The frequency cost penalized deviations from a preferred baseline frequency $f_0$, while allowing deviations above $f_0$ to be more costly than deviations below $f_0$ (*10*).

$$C_{f}(f) = \left[1 + \frac{a_{f}}{1 + exp\left[-\left(\frac{\left(f - f_{0}\right)}{s_{f}}\right)\right]}\right]\left(f - f_{0}\right)^{2}$$

where $\alpha_f$ controls the asymmetry of the cost, and $s_f$ controls the smoothness of the transition around $f_0$.

*Utility function*

The total utility of a call was defined as

$$U(A, f, d, \phi) = I(A, f, d, \phi) - C(A, f, d)$$

This objective formalizes the trade-off between the information gained from a call and the energetic costs associated with vocal production.

*Stochastic sampling of optimal call parameters*

We evaluated the utility function on a four-dimensional grid spanning call amplitude, duration, peak frequency, and phase. The grid is sampled with 121 values per parameter, amounting to a total of 104,060,401 parameter combinations. The grids were defined as follows:

$$A \in [60, 120]\ dB\ SPL$$

$$d \in [0.5, 3.0]\ ms$$

$$f \in [70, 110]\ kHz$$

$$\phi \in [0, 2\pi]$$

To generate the model predictions over call parameters, we converted the utility function into a probability distribution over the entire grid using a Softmax function:

$$P(A, f, d, \phi \mid c) = \frac{exp\left(\frac{U(A, f, d, \phi)}{T}\right)}{\sum_{A, f, d, \phi} exp\left(\frac{U(A, f, d, \phi)}{T}\right)}$$

where $T$ is a temperature parameter controlling variability. For each condition, we sampled N = 1000 combinations of call parameters from the resulting distribution. The fixed model parameters (see Table S1) were selected to place the model predictions within the empirical range of call amplitudes, durations, and frequencies. The parameters were not fitted to individual animals. The model predicts the same qualitative differences between the three conditions for other sets of parameters.

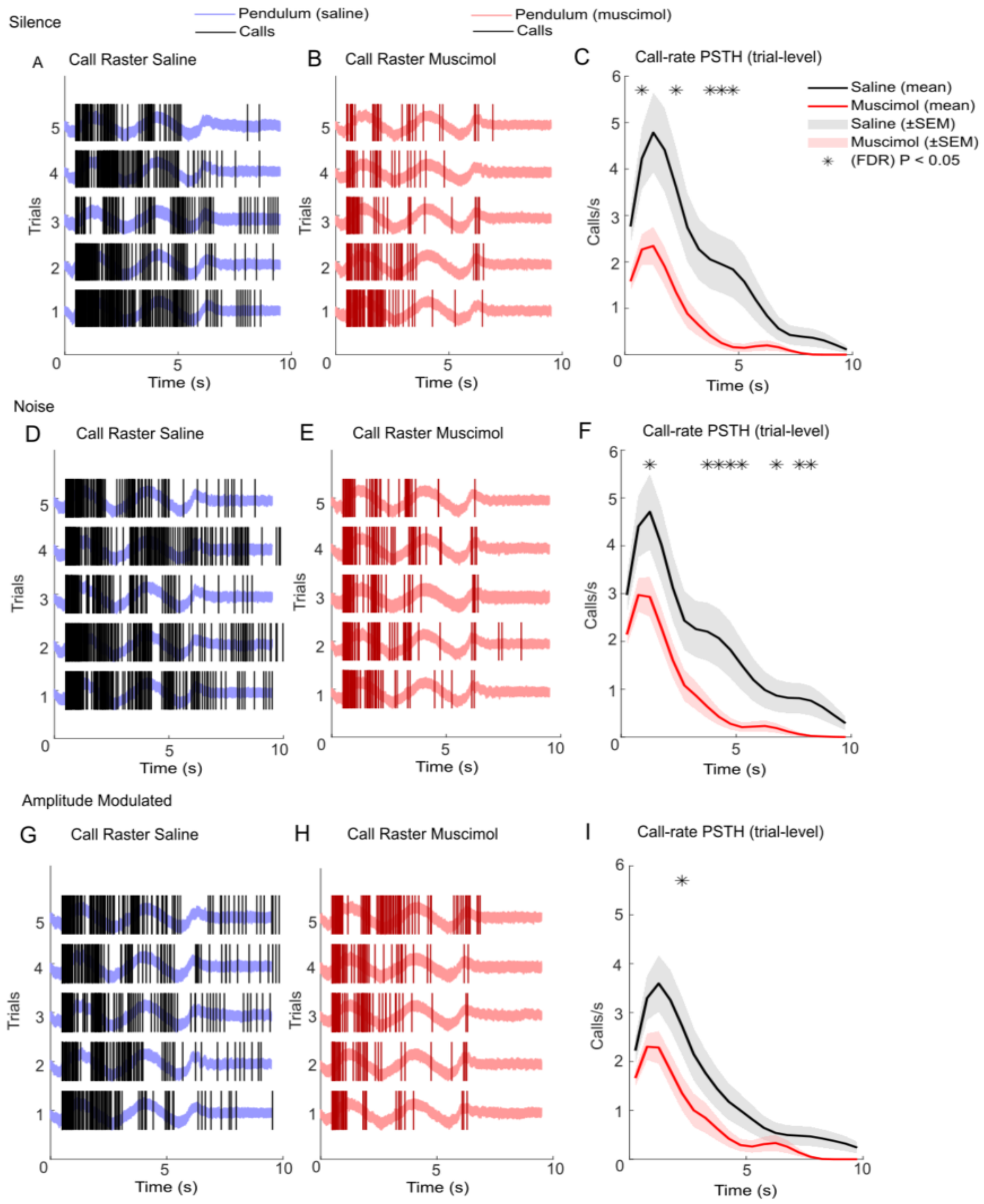


**Fig. S1. FAF inactivation suppresses call-rate dynamics during pendulum echolocation.**
(A–I) Trial-by-trial call rasters, population call-rate histograms in silence, continuous broadband noise, amplitude-modulated noise under saline, muscimol conditions. (A, B, D, E, G, H) Example rasters show pendulum position, individual calls across repeated trials. (C, F, I) Peristimulus time histograms (PSTHs) show mean call rate across trials; shaded regions denote ± SEM. Time bins with significant saline-versus-muscimol differences after Benjamini–Hochberg FDR correction (P < 0.05) indicated with (*).

**Table S1. Parameters used for the normative model**

| Parameter | Value | Description |
|---|---|---|
| $\sigma 2$ | 0.1 | Baseline noise level |
| $\beta$ | 2.0 | Sigmoid slope |
| $\theta$ | 4.0 | Sigmoid inflection point |
| $A_{ref}$ | 90 dB | SPL Baseline amplitude |
| $f_0$ | 90 kHz | Baseline frequency |
| $\alpha_f$ | 3.0 | Frequency cost asymmetry |
| $s_f$ | 0.5 | Frequency cost smoothness |
| $\lambda_f$ | $1 \times 10^{-5}$ | Frequency cost coefficient |
| $\lambda_A$ | $1 \times 10^{-4}$ | Amplitude cost coefficient |
| $\lambda_d$ | $1 \times 10^{-3}$ | Duration cost coefficient |
| $\lambda_c$ | $1 \times 10^{-3}$ | Baseline cost |
| $f_n$ | 90.0 kHz | Noise center |
| $\sigma_f^{\ 2}$ | 10.0 | Noise bandwidth |
| $N_B$ | 20.0 | Noise magnitude |
| $N_{min}$ | 0.5 | AM noise minimum |
| $N_{max}$ | 1.0 | AM noise maximum |
| $\phi_m$ | 0.0 | AM noise phase |
| $T$ | $1 \times 10^{-4}$ | Softmax temperature |

**Table S2a. Statistical results corresponding Figure 2A, C, E.**

The table reports the exact p-values and Cliff's delta (d) effect sizes for the example bat. Comparisons include Silence, Cont. Noise, and 4Hz AM noise for call intensity, duration, and peak frequency.

| Metric | Comparison | *P* value | Cliff's delta |
|---|---|---|---|
| **Intensity (dB)** | Silence vs Cont. Noise | $4.47 \times 10^{-21}$ | 0.86 |
| | Cont. Noise vs 4Hz AM Noise | 0.362 | -0.07 |
| | Silence vs 4Hz AM Noise | $3.493\times 10^{-11}$ | 0.67 |
| **Duration (ms)** | Silence vs Cont. Noise | $9.0553 \times 10^{-9}$ | 0.51 |
| | Cont. Noise vs 4Hz AM Noise | $4.755 \times 10^{-6}$ | -0.34 |
| | Silence vs 4Hz AM Noise | 0.0054 | 0.27 |
| **Peak Frequency (kHz)** | Silence vs Cont. Noise | 0.2478 | -0.11 |
| | Cont. Noise vs 4Hz AM Noise | 0.0105 | -0.2 |
| | Silence vs 4Hz AM Noise | 0.0043 | -0.29 |

**Table S2b. Statistical results corresponding Figure 2B, D, F.**

The table reports the exact p-values and Cliff's delta (d) effect sizes for the population analysis. Comparisons include Silence, Cont. Noise, and 4Hz AM Noise for call intensity, duration, and peak frequency.

| Metric | Comparison | *P* value | Cliff's delta |
|---|---|---|---|
| **Intensity (dB)** | Silence vs Cont. Noise | 0.00012 | 0.55 |
| | Cont. Noise vs 4Hz AM Noise | 0.0295 | -0.21 |
| | Silence vs 4Hz AM Noise | 0.00012 | 0.54 |
| **Duration (ms)** | Silence vs Cont. Noise | 0.00012 | 0.76 |
| | Cont. Noise vs 4Hz AM Noise | 0.00292 | -0.33 |
| | Silence vs 4Hz AM Noise | 0.00195 | 0.56 |
| **Peak Frequency (kHz)** | Silence vs Cont. Noise | 0.00012 | -0.62 |
| | Cont. Noise vs 4Hz AM Noise | 0.00024 | -0.68 |
| | Silence vs 4Hz AM Noise | 0.00012 | -0.92 |

**Table S3a. Statistical results corresponding Figure 5A, C, E.**

The table reports the exact p-values and Cliff's delta (d) effect sizes for the example bat. Comparisons include saline versus muscimol within each call type (Silence, Cont. Noise, and 4Hz AM noise) and pairwise comparisons between call types within the saline and muscimol conditions for call intensity, duration, and peak frequency.

| Metric | Comparison | *P* value | Cliff's delta |
|---|---|---|---|
| **Intensity (dB)** | Silence Saline vs Silence Muscimol | $5.53 \times 10^{-9}$ | 0.76 |
| | Cont. Noise Saline vs Cont. Noise Muscimol | $2.10 \times 10^{-4}$ | 0.45 |
| | 4Hz AM Noise Saline vs 4Hz AM Noise Muscimol | $5.43 \times 10^{-4}$ | 0.54 |
| | Silence Saline vs Cont. Noise Saline | 0.002 | 0.49 |
| | Cont. Noise Saline Vs 4Hz AM Noise Saline | 0.248 | 0.21 |
| | Silence Saline Vs 4Hz AM Noise Saline | $2.02 \times 10^{-4}$ | 0.69 |
| | Silence Muscimol Vs Cont. Noise Muscimol | 0.552 | 0.05 |
| | Cont. Noise Muscimol Vs 4Hz AM Noise Muscimol | 0.469 | 0.06 |
| | Silence Muscimol Vs 4Hz AM Noise Muscimol | 0.420 | 0.07 |
| **Duration (ms)** | Silence Saline Vs Silence Muscimol | $1.06 \times 10^{-6}$ | 0.62 |
| | Cont. Noise Saline Vs Cont. Noise Muscimol | $2.26 \times 10^{-8}$ | -0.67 |
| | 4Hz AM Noise Saline Vs 4Hz AM Noise Muscimol | 0.038 | 0.31 |
| | Silence Saline Vs Cont. Noise Saline | $4.28 \times 10^{-5}$ | 0.63 |
| | Cont. Noise Saline Vs 4Hz AM Noise Saline | 0.060 | -0.34 |
| | Silence Saline Vs 4Hz AM Noise Saline | 0.125 | 0.29 |
| | Silence Muscimol Vs Cont. Noise Muscimol | $1.68 \times 10^{-13}$ | -0.65 |
| | Cont. Noise Muscimol Vs 4Hz AM Noise Muscimol | $1.29 \times 10^{-10}$ | 0.53 |
| | Silence Muscimol Vs 4Hz AM Noise Muscimol | 0.020 | -0.19 |
| **Peak Frequency (kHz)** | Silence saline vs Silence muscimol | $7.23 \times 10^{-5}$ | 0.52 |
| | Cont. Noise saline vs Cont. Noise muscimol | $8.34 \times 10^{-16}$ | 0.98 |
| | 4Hz AM Noise saline vs 4Hz AM Noise muscimol | $5.42 \times 10^{-9}$ | 0.91 |

| | | | |
|---|---|---|---|
| | Silence saline vs Cont. Noise saline | $2.02 \times 10^{-7}$ | -0.8 |
| | Cont. Noise saline vs 4Hz AM Noise saline | 0.261 | -0.2 |
| | Silence saline vs 4Hz AM Noise saline | $3.77 \times 10^{-7}$ | -0.94 |
| | Silence muscimol vs Cont. Noise muscimol | 0.087 | 0.16 |
| | Cont. Noise muscimol vs 4Hz AM Noise muscimol | $5.06 \times 10^{-10}$ | -0.52 |
| | Silence muscimol vs 4Hz AM Noise muscimol | 0.005 | -0.24 |

**Table S3b. Statistical results corresponding Figure 5B, D, F.**

The table reports the exact p-values and Cliff's delta (d) effect sizes for the population analysis. Comparisons include saline versus muscimol within each call type (Silence, Cont. Noise, and 4Hz AM Noise) and pairwise comparisons between call types within the saline and muscimol conditions for call intensity, duration, and peak frequency.

| Metric | Comparison | *P* value | Cliff's delta |
|---|---|---|---|
| **Intensity (dB)** | Silence saline vs Silence muscimol | 0.008 | 0.81 |
| | Cont. Noise saline vs Cont. Noise muscimol | 0.016 | 0.89 |
| | 4Hz AM Noise saline vs 4Hz AM Noise muscimol | 0.250 | 0.5 |
| | Silence saline vs Cont. Noise saline | 0.641 | 0.156 |
| | Cont. Noise saline vs 4Hz AM Noise saline | 0.016 | 0.875 |
| | Silence saline vs 4Hz AM Noise saline | 0.016 | 0.969 |
| | Silence muscimol vs Cont. Noise muscimol | 0.611 | -0.018 |
| | Cont. Noise muscimol vs 4Hz AM Noise muscimol | 0.021 | 0.16 |
| | Silence muscimol vs 4Hz AM Noise muscimol | 0.009 | 0.222 |
| **Duration (ms)** | Silence saline vs Silence muscimol | 0.008 | 0.797 |
| | Cont. Noise saline vs Cont. Noise muscimol | 0.008 | -0.953 |
| | 4Hz AM Noise saline vs 4Hz AM Noise muscimol | 0.008 | 0.891 |
| | Silence saline vs Cont. Noise saline | 0.016 | 0.891 |
| | Cont. Noise saline vs 4Hz AM Noise saline | 0.016 | -0.938 |
| | Silence saline vs 4Hz AM Noise saline | 0.469 | -0.266 |
| | Silence muscimol vs Cont. Noise muscimol | 0.008 | -0.875 |
| | Cont. Noise muscimol vs 4Hz AM Noise muscimol | 0.016 | 0.891 |
| | Silence muscimol vs 4Hz AM Noise muscimol | 0.938 | 0.141 |
| **Peak Frequency (kHz)** | Silence saline vs Silence muscimol | 0.008 | 0.844 |
| | Cont. Noise saline vs Cont. Noise muscimol | 0.008 | 0.922 |
| | 4Hz AM Noise saline vs 4Hz AM Noise muscimol | 0.008 | 0.938 |
| | Silence saline vs Cont. Noise saline | 0.008 | -0.672 |

| | | | |
|---|---|---|---|
| | Cont. Noise saline vs 4Hz AM Noise saline | 0.125 | -0.141 |
| | Silence saline vs 4Hz AM Noise saline | 0.008 | -0.938 |
| | Silence muscimol vs Cont. Noise muscimol | 0.742 | -0.219 |
| | Cont. Noise muscimol vs 4Hz AM Noise muscimol | 0.078 | -0.281 |
| | Silence muscimol vs 4Hz AM Noise muscimol | 0.055 | -0.516 |

**Table S4. Statistical results corresponding Figure 5L.**

The table reports the exact *p*-values for the Rayleigh tests (saline and FAF inactivation), the preferred phase (mean angle) for each condition, and the Watson–Williams test *p*-values comparing preferred phases between conditions for each bat.

| Bat | Rayleigh p saline | Rayleigh p muscimol | Angular shift saline | Angular shift muscimol | Watson Williams p |
|---|---|---|---|---|---|
| **1** | $4.66 \times 10^{-17}$ | $7.84 \times 10^{-24}$ | -2° (CW) | +175° (CCW) | $3.18 \times 10^{-46}$ |
| **2** | $1.48 \times 10^{-20}$ | 0.206 | +30° (CCW) | -102° (CW) | $8.37 \times 10^{-8}$ |
| **3** | $4.46 \times 10^{-36}$ | 0.309 | +9° (CCW) | +178° (CCW) | $7.05 \times 10^{-13}$ |
| **4** | 0.001 | 0.188 | +39° (CCW) | -133° (CW) | $3.37 \times 10^{-7}$ |
| **5** | $4.38 \times 10^{-6}$ | 0.071 | +4° (CCW) | -153° (CW) | $1.84 \times 10^{-14}$ |
| **6** | 0.002 | $5.81 \times 10^{-4}$ | +6° (CCW) | +165° (CCW) | $1.71 \times 10^{-12}$ |
| **7** | 0.010 | 0.510 | +31° (CCW) | +67° (CCW) | 0.139 |
| **8** | $2.47 \times 10^{-4}$ | $9.74 \times 10^{-16}$ | +50° (CCW) | +145° (CCW) | $1.71 \times 10^{-8}$ |

References

1. A. Kiai, J. Clemens, M. Kössl, D. Poeppel, J. Hechavarría, Flexible control of vocal timing in Carollia perspicillata bats enables escape from acoustic interference. *Commun Biol* **6**, 1153 (2023).

2. L. M. Martin, F. García-Rosales, M. J. Beetz, J. C. Hechavarría, Processing of temporally patterned sounds in the auditory cortex of Seba's short-tailed bat,Carollia perspicillata. *European Journal of Neuroscience* **46**, 2365–2379 (2017).

3. H.-U. Schnitzler, E. K. V. Kalko, Echolocation by Insect-Eating Bats: We define four distinct functional groups of bats and find differences in signal structure that correlate with the typical echolocation tasks faced by each group. *BioScience* **51**, 557–569 (2001).

4. M. Knörnschild, K. Jung, M. Nagy, M. Metz, E. Kalko, Bat echolocation calls facilitate social communication. *Proc. R. Soc. B.* **279**, 4827–4835 (2012).

5. S. A. Zollinger, H. Brumm, The Lombard effect. *Curr Biol* **21**, R614-615 (2011).

6. C. Chiu, W. Xian, C. F. Moss, Adaptive echolocation behavior in bats for the analysis of auditory scenes. *J Exp Biol* **212**, 1392–1404 (2009).

7. M. Taub, A. Goldshtein, A. Boonman, O. Eitan, E. Hurme, S. Greif, Y. Yovel, What determines the information update rate in echolocating bats. *Commun Biol* **6**, 1187 (2023).

8. J. Luo, H. R. Goerlitz, H. Brumm, L. Wiegrebe, Linking the sender to the receiver: vocal adjustments by bats to maintain signal detection in noise. *Sci Rep* **5**, 18556 (2015).

9. I. Foskolos, M. Bjerre Pedersen, K. Beedholm, A. S. Uebel, J. Macaulay, L. Stidsholt, S. Brinkløv, P. T. Madsen, Echolocating Daubenton's bats are resilient to broadband, ultrasonic masking noise during active target approaches. *J Exp Biol* **225**, jeb242957 (2022).

10. D. K. N. Dechmann, M. Wikelski, H. J. van Noordwijk, C. C. Voigt, S. L. Voigt-Heucke, Metabolic costs of bat echolocation in a non-foraging context support a role in communication. *Front. Physiol.* **4** (2013).
